# A bacterial display system for effective selection of protein-biotin ligase BirA variants with novel peptide specificity

**DOI:** 10.1101/367730

**Authors:** Jeff Granhøj, Henrik Dimke, Per Svenningsen

**Author notes:** Corresponding author: Dr. Per Svenningsen, Institute of Molecular Medicine, University of Southern Denmark, J.B. Winsloews vej 21.3, DK-5000 Odense C, Denmark.

## Abstract

Biotinylation creates a sensitive and specific tag for purification and detection of target proteins. The *E. coli* protein-biotin ligase BirA biotinylates a lysine within a synthetic biotin acceptor peptide (AP) and allow for specific tagging of proteins fused to the AP. The approach is not applicable to unmodified proteins, and we sought to develop an effective selection system that could form the basis for directed evolution of novel BirA variants with specificity towards unmodified proteins. The system was based on bacterial display of a target peptide sequence, which could be biotinylated by cytosolic BirA variants before being displayed on the surface. In a model selection, the bacterial display system accomplished >1.000.000 enrichment in a single selection step. A randomly mutated BirA library was used to identify novel variants. Bacteria displaying peptide sequences from 13 out of 14 tested proteins were strongly enriched after 3–5 selection rounds. Moreover, a clone selected for biotinylation of a C-terminal peptide from red-fluorescent protein TagRFP showed site-specific biotinylation. Thus, active BirA variants with novel specificity are effectively isolated with our bacterial display system and provides a basis for the development of BirA variants for site-selective biotinylation.

## Introduction

Biotinylation of biomolecules is widely used in biomedical sciences due to the high affinity binding of biotin to streptavidin and its homologues, and biotin therefore creates a simple and effective tag for purification and detection of targets molecules. Proteins can be biotinylated through chemical or enzymatic techniques. The chemical techniques are simple; however, they modify a broad range of chemically similar groups and therefore often lack target selectivity. Enzymatic biotinylation, on the other hand, is highly specific and is catalyzed by protein-biotin ligases. All living organisms express protein-biotin ligases, which biotinylate between 1–5 specific proteins in their respective organisms.^1^ The *E. coli* protein-biotin ligase BirA is extremely specific and covalently attaches biotin to a single lysine in the endogenously expressed BCCP subunit of acetyl-CoA carboxylase.^2^ Additionally, a small synthetic 15-amino-acid peptide is effectively biotinylated by BirA^3,4^ and fusion of the synthetic biotin acceptor peptide (AP) to a target protein creates an efficient approach for site-specific *in vivo* and *in vitro* biotinylation through co-expression or addition of BirA.^5^-^7^ Although the high specificity and activity of BirA towards the synthetic AP provide a powerful approach for specific labeling, the range of applications for BirA mediated biotinylation is restricted to samples containing AP-fused proteins. Development of novel BirA variants with specificity towards peptide sequences that are present in endogenously expressed proteins would therefore massively expand the application range wherein enzymatic biotinylation can be utilized. Directed evolution is a powerful method to evolve new protein function and involves iterations of gene mutations, isolation and amplification of gene variants with the desired function. BirA activity is readily selected and phage display technologies have been developed that allow for selection of BirA activity based on their display of biotinylated peptides.^8,9^ The strong binding of biotin to streptavidin ensures that even low abundant phages are selected, but amplification of phages requires infection a bacterial host and the phages therefore have to be eluted from the affinity resin. The elution creates a bottleneck and, as an alternative to streptavidin, low affinity monomeric avidin has been used as it allows for reversible binding of biotinylated phages and elution with biotin.^8,9^ The lower affinity of monomeric avidin affects the selection of biotinylated phages, and allowed only for a modest ~10-fold enrichment of BirA from phages expressing synthetic AP.^8^ To accelerate the identification of active BirA variants, we sought to develop a selection method in which the elution step could be eliminated.

Bacteria can be selected and enriched through magnetic beads isolation with no need for elution^10^, and we therefore based our selection on a bacterial display system. As scaffold for peptide display, we used eCPX (enhanced circularly permuted outer membrane protein OmpX), which allow for effective display of peptides at both N- and C-termini.^11,12^ The target peptide sequence is inserted into the carboxy-terminus of eCPX^12^ such that bacteria expressing active BirA variants biotinylate and subsequently display the biotinylated peptide on the surface (Figure 1a), allowing for effective streptavidin selection. Here, we demonstrate that bacterial display of a peptide sequence is an effective approach to select for BirA variants with novel specificity.

**Figure 1.**
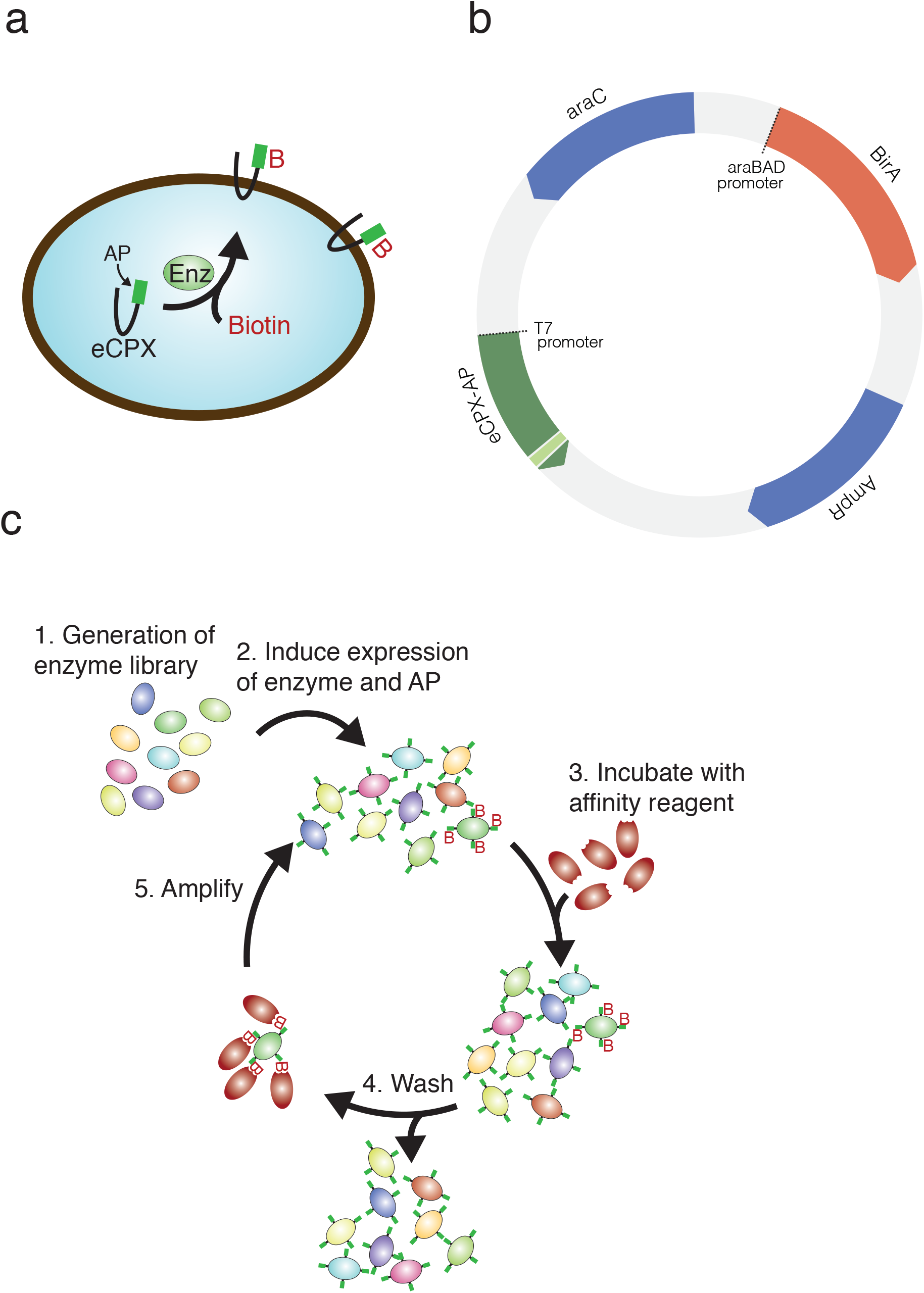
The selection system. (a) The selection system is based on bacterial surface display in which eCPX with a terminal peptide sequence is co-expressed with an enzyme (Enz). eCPX is translocated to the bacterial cell surface. The surface display of the biotin depends on protein-biotin ligase activity. (b) The bacterial display system is based on a single plasmid encoding the 2 components: the enzyme, BirA, under the control of an arabinose-inducible promoter and eCPX under the control of the T7 promoter. (c) Bacterial transformed with randomly mutated enzyme variants (1). Following a short induction period in order to express the enzyme and eCPX with the AP (2), the bacteria are incubated with a streptavidin beads (3) and unbound bacteria are removed by extensive washing (4). The streptavidin beads are diluted directly in medium and amplified overnight (5).

## Results

### Model selection of BirA against the synthetic AP

We generated a single plasmid in which BirA with a C-terminal 6xHis tag (BirA-6xHis) was controlled by an arabinose-inducible promoter (araBAD) and eCPX expression was controlled by a T7 promoter, allowing for induction of BirA-6xHis and eCPX expression using arabinose and isopropyl β-D-1-thiogalactopyranoside (IPTG), respectively (Figure 1b). The principle of the selection system is that a library of BirA mutants is co-expressed with eCPX fused to target peptide sequence and bacteria displaying biotinylated target peptide are readily selected and amplified while bound to the streptavidin beads (Figure 1c). We carried out a model selection using the BirA against its synthetic 15-amino-acid AP or its non-biotinylatable K10A sequence (AP(K10A))^7^. Biotinylated eCPX-AP was detected in uninduced cultures (Figure 2a), indicating a low basal expression of the eCPX-AP from the T7 promoter even in the absence of IPTG and sequent biotinylation by endogenous BirA. BirA-6xHis was detected in cultures of eCPX-AP and –AP(K10A) after co-induction with arabinose and IPTG, but strong biotinylation was only detected in the eCPX-AP bacteria and not in BirA-eCPX-AP(K10A) (Figure 2a).

**Figure 2.**
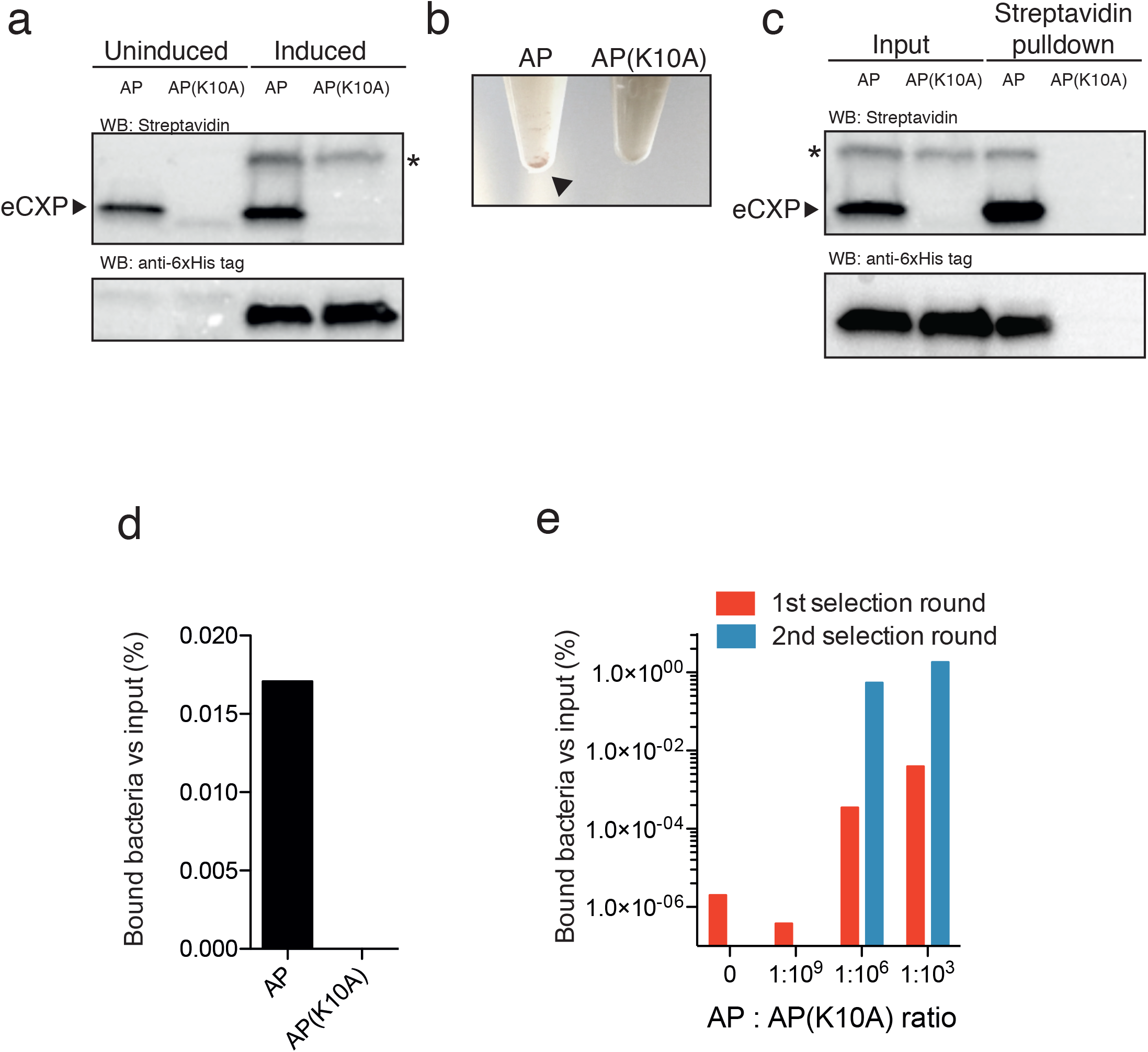
Effective enrichment of surfaced displayed biotinylated AP. (a) Western blots using streptavidin showed that eCPX-AP was biotinylated in uninduced and induced bacteria, while eCPX-AP(K10A) was not biotinylated even after induction of BirA (Histag). Uncropped versions of the blots are shown in Supplementary Figure 1. * indicates an unspecific streptavidin-reacting protein when BirA is induced. (b) Incubation of magnetic streptavidin-Dynabeads with eCPX-AP bacteria caused rapid precipitation of the beads (arrow), while no precipitate was evident in AP(K10A) bacteria. (c) The precipitate from the streptavidin pulldown of the bacteria contained BirA-6xHis in bacteria expressing AP, but not AP(K10A), while BirA-6xHis was present in input. Uncropped versions of the blots are shown in Supplementary Figure 1. (d) The streptavidin pulldown precipitated viable eCPX-AP expressing bacteria. (e) A model selection of different eCPX-AP:AP(K10A) ratios showed that bacteria were isolated and effectively enriched in cultures with eCPX-AP:AP(K10A) ratios of 1:10^3^ and 1:10^6^ after 2 selection rounds, while no enrichment was observed in cultures with eCPX-AP:AP(K10A) ratios of 0 and 1:10^9^.

The IPTG- and arabinose-induced bacteria were incubated with streptavidin-Dynabeads. Immediately upon addition of streptavidin-Dynabeads to cultures, aggregation and precipitation of the streptavidin-Dynabeads was observed in the culture with eCPX-AP, but not in eCPX-AP(K10A) (Figure 2b). Western blot of the isolated streptavidin-Dynabeads incubated with bacteria displaying the synthetic AP showed intense biotinylation of eCPX-AP as well as isolation of BirA-6xHis, while both proteins were undetected in western blots of the streptavidin pull-down from eCPX-AP(K10A) (Figure 2c). In agreement, streptavidin precipitated viable bacteria to a higher degree in eCPX-AP cultures compared to eCPX-AP(K10A) (Figure 2d). Thus, the results suggest that bacteria display biotinylated eCPX on surface and allows for isolation of the biotinylated AP. It is imperative that the selection procedure can isolate rare functional clones in the library and we therefore tested if eCPX-AP clones could be isolated after dilution into cultures of non-biotinylatable eCPX-AP(K10A) at ratios of 1:10^3^, 1:10^6^, and 1:10^9^. After 2 rounds of selection, a strong enrichment of bacteria was observed in the mixed cultures containing eCPX-AP diluted 1:10^3^ and 1:10^6^ with cultures of eCPX-AP(K10A) bacteria, whereas no enrichment was detected in the cultures consisting of 1:10^9^ dilution or cultures of pure eCPX-AP(K10A) (Figure 2e). The AP/ AP(K10A) region of the plasmid was sequenced in 5 individual clones to estimate the enrichment factor. The enrichment for eCPX-AP in the 1^st^ selection round was the highest and estimated to be in the order of 10^6^ (Table 1), suggesting that the proposed selection scheme allows for the isolation of rare variants in a BirA library.

**Table 1.** The displayed AP region was sequenced in 5 randomly selected clones from the 1^st^ and 2^nd^ selection round from the cultures mixed at a AP:AP(K10A) ratio of 1:10^3^ and 1:10^6^. The enrichment was calculated as [output AP:AP(K10A) ratio]/ [input AP:AP(K10A) ratio].

| Initial ratio<br>AP:AP(K10A) | After 1st selection |  |  | After 2nd selection |  |  |
| --- | --- | --- | --- | --- | --- | --- |
|  | AP | AP(K10A) | Enrichment | AP | AP(K10A) | Enrichment |
| **1:10<sup>3</sup>** | 5 | 0 | 5×10<sup>3</sup> | 5 | 0 | 1 |
| **1:10<sup>6</sup>** | 4 | 1 | 4×10<sup>6</sup> | 5 | 0 | 20 |

### Selection of BirA variants biotinylating novel acceptor peptides

We tested if the selection scheme allowed for isolation of BirA variants that biotinylate peptide sequences present in unmodified proteins. Lysine-containing peptide sequences from 4 different subunits from membrane proteins were tested: Na^+^/K^+^-ATPase α1 (Figure 3a), αENaC (Figure 3b), βENaC (Figure 3c) and γENaC (Figure 3d) subunits. The peptide sequences were fused to the C-terminal of eCPX and co-expressed with a BirA library. A clear enrichment in streptavidin binding bacteria were detected after 4–5 rounds of selection (Figure 3a-d). After the last selection round, 10 randomly selected clones were tested by western blotting for their ability to biotinylate the displayed peptide sequences. Most of the tested clones showed a streptavidin-reacting band migrating in parallel with the positive control expressing BirA-6xHis and eCPX-AP (Figure 3a-d). The intensity of the band was lower than that of the positive control, indicated that the isolated clones had lower activity towards the peptide sequence than BirA-6xHis had towards the synthetic AP. Moreover, we observed additional streptavidin-reacting bands with a higher molecular weight in some isolated clones expressing peptide sequences derived from Na^+^/K^+^-ATPase (Figure 3a) and γENaC (Figure 3d), indicating the isolated BirA variants had a different specificity. We tested an additional 9 peptides sequences derived from different proteins. Bacteria displaying a peptide sequences from enhanced green fluorescent protein (EGFP) did not produce an enrichment even after 4 selection rounds (Figure 4a); however, a strong enrichment was detected for the remaining displayed peptide sequences after 3–5 rounds of selection (Figure 4b-i). Thus, bacterial display systems allow for selection of BirAs with novel specificity.

**Figure 3.**
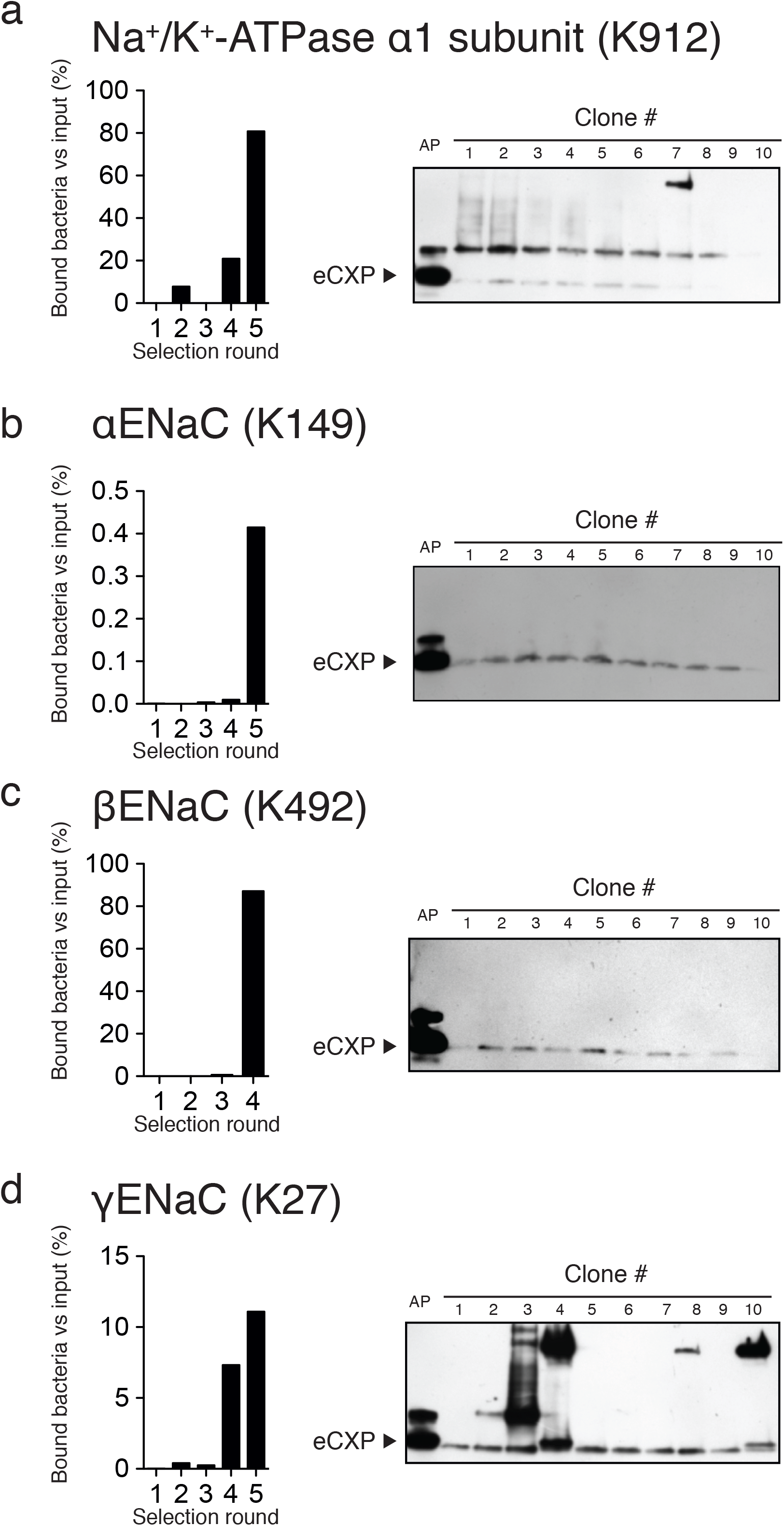
Selection of bacteria expressing BirA variants that biotinylates peptides derived from Na^+^ transporters. Bacteria displaying a peptide from (a) Na^+^/K^+^-ATPase α1 subunit, (b) αENaC, (c) βENaC and (d) γENaC were isolated through 4–5 selection rounds, and biotinylation of the eCPX-displayed peptide was tested by western blotting. Bacteria displaying the synthetic AP was included as positive controls. The majority of the isolated clones produced a band consistent with biotinylation of the displayed peptide. The intensity of the band was, however, low compared to AP, and in some of the clones additional streptavidin reacting bands were present at higher molecular weights. Uncropped versions of the blots are shown in Supplementary Figure 2.

**Figure 4.**
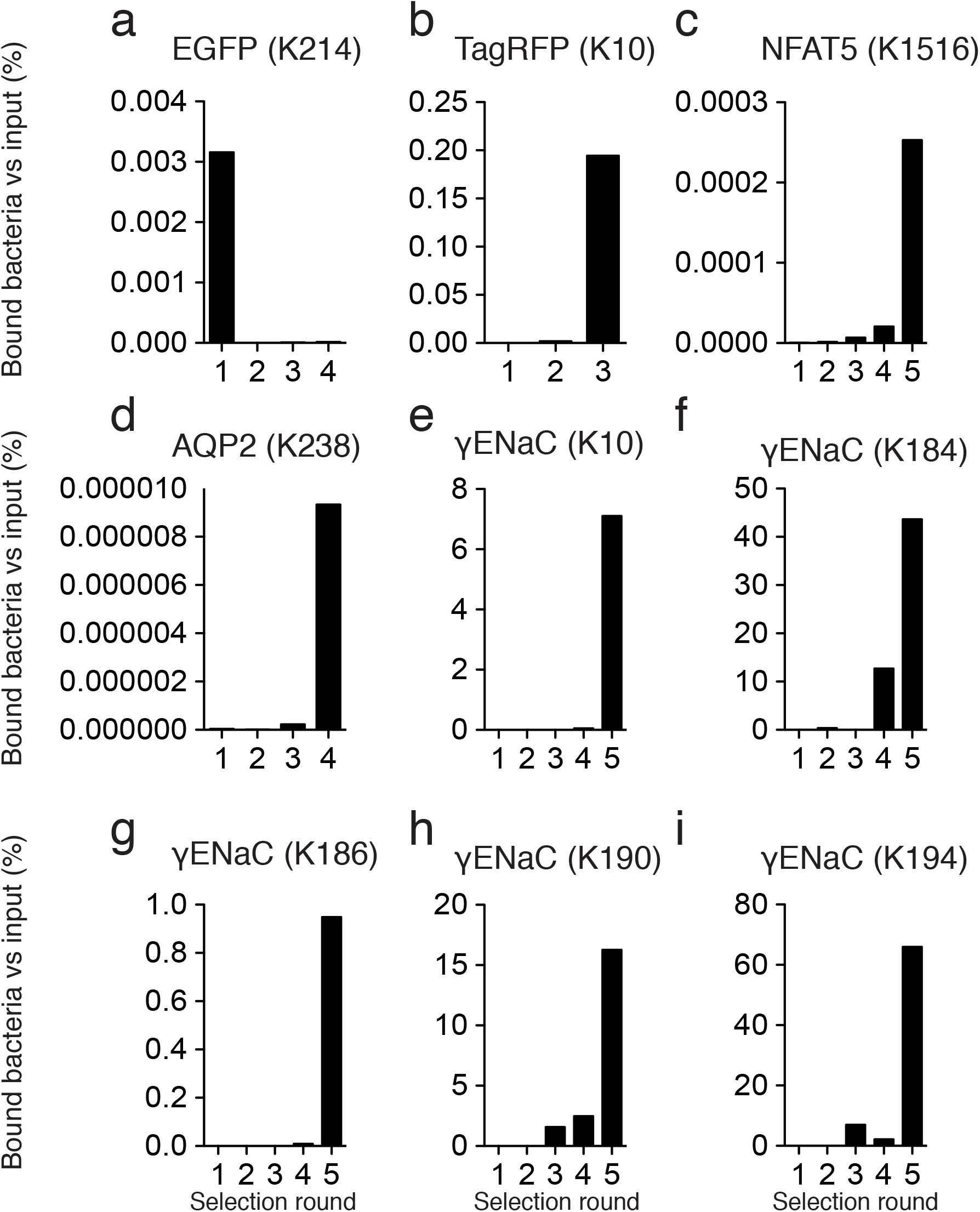
Selection of bacteria expressing BirA variants with novel activity. Bacteria displaying peptides derived from (a)EGFP, (b) N terminal of TagRFP, i.e. TagRFP(K10),(c)NFAT5, (d) AQP2, (e) γENaC subunit K10, (f) K184, (g) K186, (h) K190 and (i) K194. Through 3–5 selection rounds, a clear enrichment of streptavidin binding bacteria were observed for all displayed peptides (b-i), except (a) EGFP.

### Biotinylation of TagRFP

We next tested if BirA variants selected for biotinylation of specific peptide sequence, could also biotinylate the native protein. We used a peptide sequence derived from the C-terminal of red fluorescent protein TagRFP that contains 2 lysines: K231 and K235 (Figure 5a). BirA-TagRFP clones were readily isolated after 3 rounds of selection (Figure 5b), and 10 clones were randomly selected to further characterize their ability to biotinylate TagRFP by co-transforming them with a plasmid encoding TagRFP fused to the C-terminal of maltose binding protein (MBP). In 8 of the 10 clones, MBP-TagRFP and BirA-TagRFP co-expression biotinylated a protein migrating with a band size of ~75kDa, consistent with the expected molecular weight of MBP-TagRFP (Figure 4c). In clone 3 and 7, the biotinylated band at ~75 kDa was not detectable; however, a faint band at the expected molecular weight of eCPX (~22 kDa) was observed. Clone 1 was used to test the specificity of the TagRFP biotinylation. Lysates from cell expression BirA-6xHis or BirA:TagRFP Clone 1 were mixed with MBP-TagRFP or MBP-TagRFP with K231A, K235A and K231A,K235A mutations. MBP was readily detected in all the tested combinations of lysates (Figure 5d). Expression of BirA-6xHis caused a strong biotinylation of eCPX-AP, but no biotinylation product was detected at the expected size of MBP-TagRFP (Figure 5d). In contrast, biotinylated bands at 75 kDa were detected after incubation of MBP-TagRFP and MBP-TagRFPK235 lysates with Clone 1, and the intensity of this band was strongly reduced by a mutation of K231A and K231A,235A in TagRFP (Figure 5d). Similar to the clones isolated from bacteria expressing peptides from Na^+^/K^+^ - ATPase α1 subunit, as well as the αENaC, βENaC and γENaC subunits, the intensity of the 75 kDa bands were low compared to the intensity of the eCPX-AP band from the BirA-6xHis expressing bacteria. The results, however, indicate that BirA selected with the bacterial display system allows for biotinylation of a specific protein in complex mixtures.

**Figure 5.**
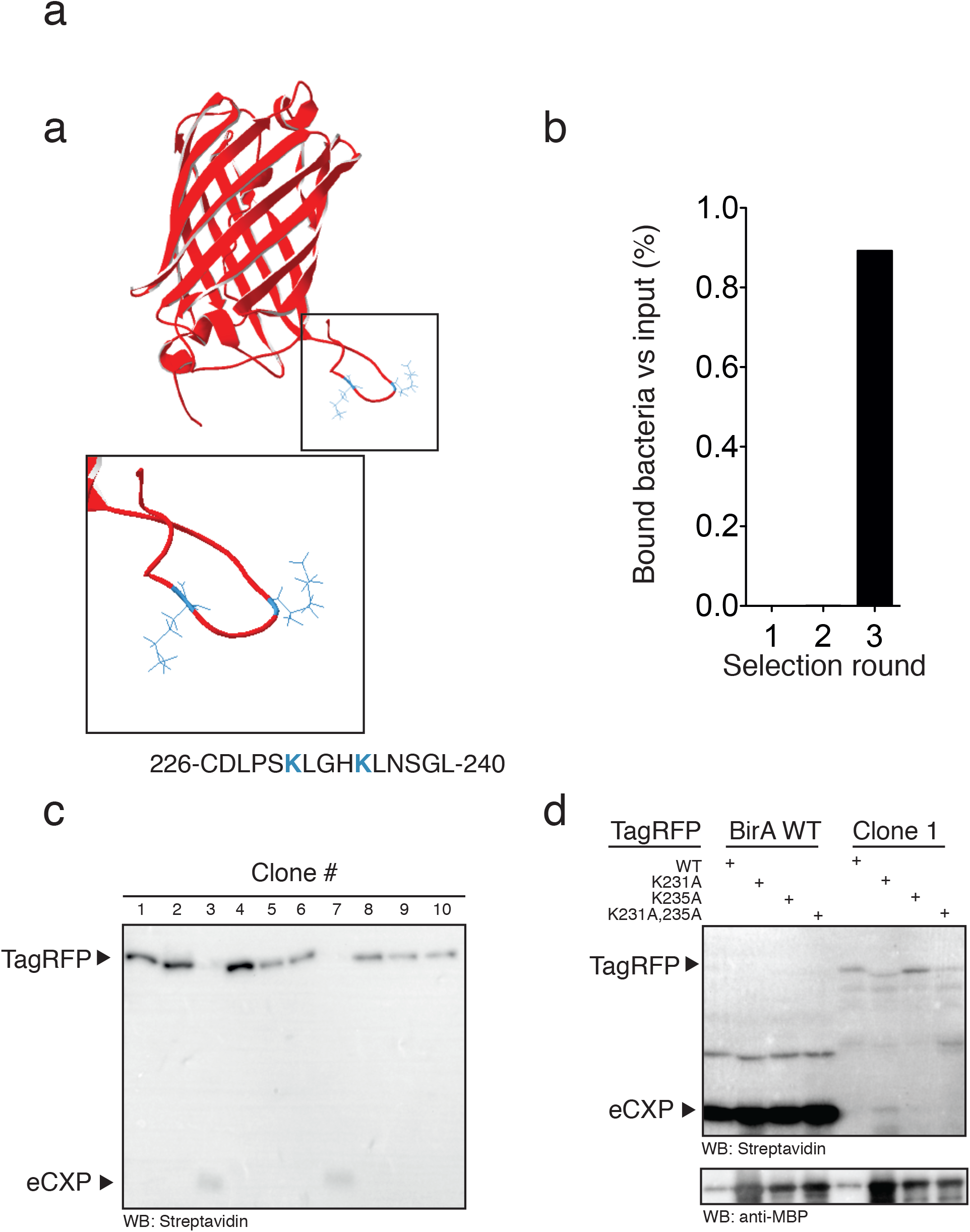
Biotinylation of TagRFP. (a) The C-terminal of TagRFP has 2 lysines (blue) at residue 231 and 235. (b) Bacteria displaying the C-terminal peptide from TagRFP were clearly enriched at 3 selection rounds using streptavidin Dynabeads. (c) Ten clones isolated after 3 selection rounds were transformed with MBP-TagRFP (expected size 75 kDa). Western blots of the clones showed that 8/10 clones caused the appearance of a streptavidin reacting band at 75 kDa, whereas the 2 negative clones showed a faint band at 22 kDa, which is the expected size of eCPX. An uncropped version of the blot is shown in Supplementary Figure 3. (d) MBP-TagRFP was not biotinylated by BirA-6xHis, but clone 1 biotinylated TagRFP and TagRFP with K235A mutation. Western blot against MBP showed that TagRFP was present in all lanes. Uncropped versions of the blots are shown in Supplementary Figure 3.

### Potential targets

Specific protein biotinylation could provide a novel approach for purification and imaging of target proteins in complex mixtures, and we analyzed a total number of 16966 mouse proteins and 20316 human proteins for their lysine abundance. Only 54 mouse proteins (0.3%) and 111 human proteins (0.5%) did not contain lysine in their primary sequence, and the majority of mouse and human proteins contained 1 or more lysine(s) per protein (Figure 6a). Since structural restrains might limit BirA’s ability to biotinylate its target protein, we restricted our analysis to the initial and terminal 30 N- and C-terminal amino acid residues of each protein. In the mouse and human proteomes, 76.6% (12989 proteins) and 75.8% (15404 proteins) of the proteins, respectively, had one or more lysine(s) in their first and/or last 30 amino acid residues (Figure 6b and c, Supplementary Data 1). This indicates that directed evolution of BirA could potentially be used to produce a broad range BirA variants that biotinylates specific proteins in complex mixtures.

**Figure 6:**
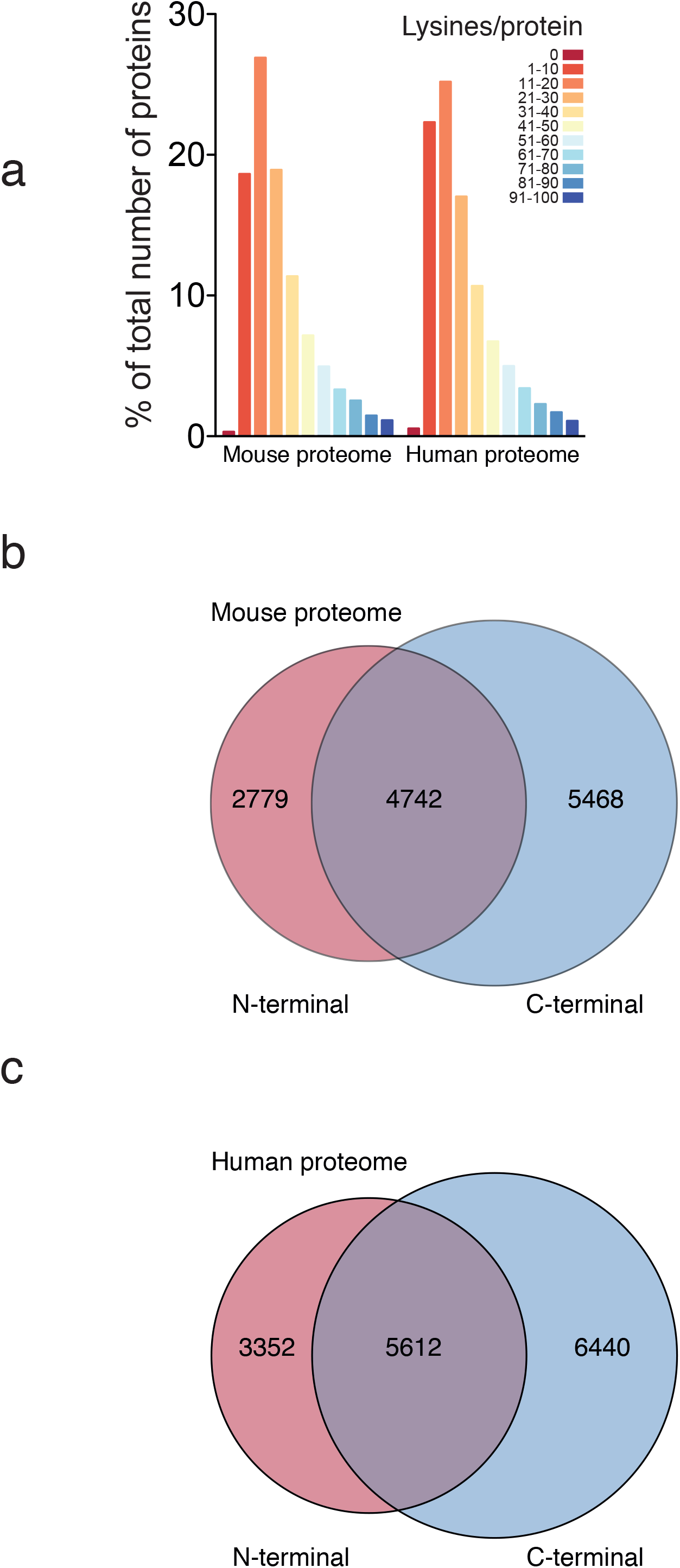
Bioinformatic analysis of lysines in the mouse and human proteome. (a) 16966 mouse proteins and 20316 human proteins were analyzed for their lysine abundance. (b) The analysis for abundance of lysines were restricted to the initial and last 30 amino acid residues for the mouse proteins, and 4742 proteins had both N- and C-terminal lysines, while 2779 and 5468 had lysines present in the N or C-terminal, respectively. (c) The analysis for abundance of lysines were restricted to the initial and last 30 amino acid residues for the human proteins, and 5612 proteins had both N- and C-terminal lysines, while 3352 and 6440 had lysines present in the N or C-terminal, respectively.

## Discussion

Using a bacterial display system, we have demonstrated that bacteria expressing BirA variants can be selected for their ability to biotinylate specific peptide sequences and that the system allows for strong enrichment of active clones by streptavidin pulldown and direct amplification of the streptavidin-bound bacteria. The bacterial display system, thus, provides a basis for directed evolution of BirA variants with activity targeted towards specific target proteins. Almost all human and mouse proteins contain lysines and novel BirA variants could provide an attractive means for site-specific protein biotinylation in complex mixtures and thereby provide a high affinity tag for downstream purification or imaging of specific proteins of interest.

The bacterial selection system effectively isolated BirA-6xHis activity against the synthetic AP diluted 10^6^-fold in cultures of bacteria displaying a mutated AP. The enrichment is orders of magnitude higher than previously reported for the phage display systems using monomeric avidin selection,^8^ indicating that the higher affinity of streptavidin increase the stringency of the selection process. Streptavidin selection of phages displaying biotinylated peptides has been used to identify novel peptide substrates for yeast biotin protein ligase, and yielded an estimated enrichment factor of ~2000.^13^ The streptavidin-bound phages were eluted by incubation at high temperatures,^13^ indicating that the strong biotin-streptavidin binding combined with the multivalency of the displayed peptide (~5 peptides are displayed per phage^14^) ensure isolation of rare active variants, but the high avidity still poses a problem in that incomplete elution of the isolated phages could hamper enrichment. The bacterial display system may, thus, have an advantage in that the elution can be eliminated and therefore allowed us to combine high affinity selection with amplification. The bacterial display system allowed for the isolation of clones with novel peptide substrate specificity. Compared to the BirA biotinylation of the synthetic AP,^3^ however, our isolated clones showed relatively low activity towards the displayed peptide sequences. For directed evolution of our isolated BirA variants, it will be important to select for biotinylation rate by increasing selection presure through e.g. lowered expression of BirA or shorter reaction time. In addition to reaction rate, the target specificity of the selected BirA variants are important. We observed that some clones isolated from the display of peptides from Na^+^/K^+^-ATPase α1 subunit and γENaC (K27) biotinylated additional bands. Promiscuous BirA activity is, however, readily detected by western blotting and the specific and promiscuous BirA variants could be used as precursors for novel BirA variants, which through e.g. DNA shuffling^15^ could be used to evolve BirA variants that biotinylates native proteins *in vivo* in e.g. mammalian cells.

In addition to its use as an affinity tag, site-specific biotinylation could potentially be used as organelle-specific *in vivo* inhibitors of post-translational modifications. Proteolytic cleavage at specific sites in the α- and γ-subunits of the epithelial sodium channel (ENaC) activates the channel.^16^-^18^ We have previously developed a monoclonal antibody directed against the neo-epitope created after proteolytic cleavage at a site corresponding lysine-186 of γENaC;^19^ however, the antibody does not allow for testing the functional consequences of organelle-specific inhibition of cleavage. We isolated BirA variants against lysine 184, 186 and 190, which could serve as precursors for development of highly activate variants that could be targeted to the lumen of the endoplasmic reticulum, Golgi apparatus and endosomes and establish in which compartment the activation occurs. Thus, the expression of the BirA variants in specific cellular organelles could provide novel spatially restricted inhibitors of post-translational modifications, such as proteolytic cleavage and ubiquitination. The selection system is not limited to BirA and could potentially be used to select for other enzymes that carries out post-translational modification, such as kinases and ubiquitinases, and allow for development of novel tools to dissect signaling pathways.

In summary, BirA variants with novel peptide substrate specificity are readily isolated using the bacterial display system. The isolated BirA variants can be used as a starting point for directed evolution to select for highly effective clones, providing tools for effective detection and isolation of endogenously expressed proteins of interest in complex *in vivo* and *in vitro* environments.

## Methods

### Constructs

The coding sequence for BirA was synthesized by Genescript and cloned into pBAD/TOPO ThioFusion (ThermoFischer Scientific). eCPX wa synthesized by GeneArt and cloned into pF1K T7 (Promega). The region covering T7 promotor, eCPX and T7 termination was amplified by PCR and inserted into the PCR amplified pBAD BirA plasmid. The final constructs pBAD BirA/eCPX was transformed into T7 Express (New England Biolabs). Insertion of peptide sequences into the C-terminal of eCPX was done by PCR. All constructs were verified by sequencing (Eurofins Genomics). A library of BirA mutants were generated using GeneMorph II EZClone Domain Mutagenesis Kit (Agilent Technologies) following the manufacturer’s instructions. The coding sequence for TagRFP was PCR amplified from pTagRFP-actin (Evrogen) and inserted into pETMBP by replacing mSA2 in the plasmid pET-MBP-mSA2 (a gift from Sheldon Park, Addgene plasmid # 52319). Site-directed mutagenesis of pET-MBP-TagRFP was carried out by PCRs. *Selection of BirA mutants*

BirA libraries were grown to OD_600_ 0.5 in 10 ml lysogeny broth (LB) and expression was induced with 0.2% L-arabinose (Sigma), 100 µM IPTG (Sigma) along with 100 µM biotin (Sigma). The cultures were grown for 1 hour at 37 °C/200 rpm. The cultures were pelleted at 5000 ×g, and washed once in ice-cold PBS (ThermoFischer Scientific). The cultures were resuspended in 500 µl ice-cold PBS. 10 µl was removed and used to calculate the input of bacteria. 20 µl Dynabeads MyOne Streptavidin C1 magnetic beads (ThermoFischer Scientific) were added to the 490 µl, and incubated for 30 min on ice. Next, the cultures were applied onto a MACS LS Column (Miltenyi Biotec) placed in a QuadroMACS (QuadroMACS), and washed extensively with ice-cold PBS. The columns were removed from the QuadroMACS and beads were eluted in 1 ml of ice-cold PBS. Ten µl of the eluted bacteria were used to calculate the retained bacteria, and the remaining elute was grown overnight at 37 °C/200 rpm. The selections process was repeated until a clear enrichment was detectable. For the model selection, The enrichment was calculated as [output AP:AP(K10A) ratio]/ [input AP:AP(K10A) ratio].

### Protein expression

pET-MBP-TagRFP (and the K231A, K235A and K231A,K235A variants) were transformed into T7 Express lysY/Iq (New England Biolabs) and grown in LB agar plates with kanamycin (Sigma Aldrich) as selection agents. A single colony was picked and grown overnight at 37 °C/200 rpm in LB medium supplemented with kanamycin. The overnight culture was diluted 1:100 in 10 ml of LB medium supplemented with kanamycin and grown for 2 hours at 37 °C/200 rpm. Protein expression was induced by addition of 1 mM IPTG and the cells were grown at 37 °C/200 rpm. The bacteria were pelleted and lysed in Cellytic B (Sigma Aldrich) diluted in PBS supplemented with 1 mM CaCl_2_ and 1 mM MgCl_2_ (PBS-CM). Expression of BirA in the selected clones was done by addition to the culture medium of 0.2% L-arabinose (Sigma Aldrich) and the cultures were grown at 37 °C/200 rpm.

### In vitro biotinylation

Lysates from BirA-TagRFP clone 1 and bacteria expressing TagRFP variants were mixed in a ratio of 1:1, supplemented with 3 µM biotin and incubated at room temperature.

### Western blotting

Bacterial lysates were mixed with LDS samples buffer (ThermoFischer Scientific) and Reducing agent (ThermoFischer Scientific) at heated for 5 min at 95 °C. The samples were loaded on 10% SDS-PAGE gels (Biorad) and run at 200V. The proteins were electroblotted onto PVDF membranes for 2h at 350 mA, and the membrane was subsequently blocked in 5% skim milk (Sigma Aldrich) in PBS supplemented with 0.05% Tween-20 (PBST) for 1 h. Biotinylation was detected by probing the membrane with streptavidin-HRP (Dako) diluted 1:1000 in PBST. MBP was detected with anti-MBP (1:5000, New England Biolabs) and secondary goat anti-mouse HRP conjugated antibody (Dako).

### Bioinformatics

The TagRFP structure was predicted using I-TASSER^20^ and analyzed using Swiss-PdbViewer^21^. Reviewed mouse and human protein sequences were downloaded from UniProt^22^ and analyzed for lysine content using Python 2.7.

### Data availability

All data generated or analysed during this study are included in this published article (and its Supplementary Information files).

## Acknowledgements

The authors thank Mohamed Abdullahi Ahmed for expert technician assistance. This work was supported by grants from the Lundbeck Foundation, the Novo Nordisk Foundation, the Danish Kidney Association, the Aase og Ejnar Danielsen Foundation, the A.P. Møller Foundation for the Advancement of Medical Science, and Knud and Edith Eriksen Memorial Foundation.

## Competing Financial Interests statement

None.

## Author Contributions

JG, HD and PS designed, performed and analyzed experiments. PS drafted the manuscript with input from co-authors. All authors reviewed and approved the final version of the manuscript.

